# The genome of *Pasteuria ramosa* reveals a high turnover rate of collagen-like genes

**DOI:** 10.1101/2024.02.09.579640

**Authors:** Alix Thivolle, Marjut Paljakka, Dieter Ebert, Peter D. Fields

## Abstract

Collagen-like proteins (CLP) are commonly found in many pathogenic bacteria where they serve as adhesins to attach to host tissue. The repetition of the amino-acid pattern (Gly-Xaa-Yaa)_n_ is the major feature of collagen and is essential to the formation of its stable triple helical structure. In the *Daphnia magna*–*Pasteuria ramosa* system, a model system for studying antagonistic coevolution, a specific CLP in the virulent parasite *P. ramosa* plays a pivotal role in host attachment, regulated by matching allele model. Recognizing the crucial role of CLPs in the infection process, we aimed to enhance our understanding of *P. ramosa*-CLPs by sequencing high-quality genomes of two isolates, using long-read technology. An analysis of a CLP gene tree of representative *Bacillota* species revealed a clear radiation of these genes in *P. ramosa*, which was not found in the closely related *Pasteuria penetrans*. A comparison of the isolates reveals a high synteny, with the exception of a few duplications and inversions, mainly involving CLPs or transposases. Across isolates, we observed a recent burst of transposases as well as duplications of CLP genes. On average, CLP genes are well conserved between isolates, but the presence/absence of individual CLP genes is not fully shared, with 39 and 43 genes in the two isolates. Our findings suggest a rapid radiation of CLP genes combined with a birth and death process of the large *P. ramosa*-CLP gene family, possibly driven by transposition and coevolution.

**Importance:** Although the host–pathogen *Daphnia magna*–*Pasteuria ramosa* system has served as a model for coevolution, we have, to date, lacked high-quality genomic resources for the parasite, as is the case for many such systems. By presenting a complete assembly of two distinct *P. ramosa* isolates, our study addresses this lack and provides deeper insights into the *P. ramosa* Collagen Like Protein (CLP) family, essential proteins involved in attachment to the host. We discover that the rapid radiation of CLP genes in *P. ramosa* appears to be driven by transposition and coevolution, enabling the parasite to adapt to host resistance mechanisms. These insights improve our understanding of host–parasite interactions and pave the way for comparative genomic analyses to better understand the evolution of these genes. They also have broader implications for disease control and therapeutic development targeting pathogenic bacteria adhesion mechanisms.

## Introduction

Collagen is the major structural component in extracellular matrices of metazoans from sponges to humans (1) and the most abundant protein in animal tissues. What distinguishes collagen from other proteins is the repetition of the Gly-Xaa-Yaa pattern necessary for forming its stable triple helical structure, giving strength to metazoan tissues. In 1998, the first (Gly-Xaa-Yaa)_n_ repeat was discovered in the bacterium *Klebsiella pneumoniae* (2), followed by further collagen-like proteins (CLPs) found in the human-pathogenic species *Streptococcus pyogenes* (3, 4), *Bacillus anthracis* (5), and *Legionella pneumophilia* (6). Besides providing structural support to the extracellular space, collagen proteins have a variety of functions including cell signaling and host immune defenses (7, 8). They can be involved in multi-drug resistance (9), biofilm formation (10), host-pathogen interaction (11) as adhesion to host tissue (12). The latter is usually the first step in an infection process, enabling the internalization or persistence of the pathogen (13). In *P. penetrans*, the dense layer of CLPs at the surface of the spores is speculated to adhere to the microvilli on the surface of its nematode host (14). A recent study of the obligate parasite *P. ramosa* has revealed that its attachment to a specific genotype of its planktonic crustacean host is mediated by the C-terminal domain of the collagen-like protein Pcl7 (15, 16).

*P. ramosa* is an obligate and virulent parasite of water fleas (*Daphnia*). It has mainly been studied for its role in the coevolutionary process with its host, revealing patterns of Red-Queen dynamics (15, 17–20). A key step of its infection process is the attachment of its transmission stage (the spore) to the host’s cuticle (21). This attachment exhibits a very high genotypic specificity that follows a matching allele model, i.e., only specific combinations of host and parasite alleles allow attachment, and thereby infection, to occur (19, 20), suggesting that these genes are critical players for the coevolution of this system. Compared to many host–parasite systems that show a more complex, and possibly even a more diffuse co-evolutionary dynamic, research on the *Daphnia*–*Pasteuria* host–parasite system has suggested that genes related to resistance and infectivity are few in number and highly specific in their function. Although several genomic regions associated with parasite resistance in the host have been mapped (22–25), the specific genes are still unknown. In the parasite, a strong correlation between polymorphisms in a single CLP gene, *Pcl7*, and an infection phenotype was discovered (15) and later substantiated through a genetic manipulation study (16). *Pcl7* is part of *P. ramosa*’s CLP gene family, which has exceptionally high diversity-more than 50 genes (26). This contrasts strongly with most other bacteria that have only three or less CLP genes (26). The extensive size of *P. ramosa’s* CLPs family suggests that other *P. ramosa*-CLPs might also be involved in the attachment process of specific parasite strains to specific host genotypes.

The diversity of *P. ramosa* CLP genes is thought to be the mechanism that drives the highly genotypic specificity of its infection in *D. magna*, and may be the result of ancient host–pathogen coevolution (27). Consequently, the development of a high-quality reference genome assembly for *P. ramosa* becomes imperative to propel research in this model system. Understanding this specificity is also crucial for understanding *Pasteuria* spp. role as a biocontrol agent of plant-pathogenic nematodes, where collagen is also believed to be of crucial importance (14, 28). We sequenced two geographically distinct *P. ramosa* isolates (P21 from Switzerland, C1 from Russia), each with different infectotypes with reference to their host genotype. The P21 isolate was sequenced using PacBio technology, while the C1 isolate genome was assembled using a combination of long-read ONT technology and short-insert Illumina reads. We obtained a circular chromosome with, respectively, 1455 and 1426 structural annotations for these isolates, including 39 and 43 putative *P. ramosa-*CLP genes, as well as an unusually high number of transposases. A phylogenetic, tree based on the P21 genome, places *P. ramosa* firmly in the *Bacillota* phylum and reveals a monophyletic group formed with *P. penetrans* and *Thermoactinomyces sp.*. The comparison between the two genomes revealed a strong synteny except for a large inversion (∼174 kb), as well as a shorter duplication (∼6 kb), both of which involve CLP genes. Together, our findings suggest rapid evolution within the CLP gene family.

## Results

### Genome metrics

The genome assemblies of *P. ramosa* resulted in single contigs of 1.77 Mb for P21 and 1.74 Mb for C1, with 1455 and 1426 structural annotations, respectively (Figure 1A for P21, Table 1). In the P21 assembly, protein coding sequences (CDS) represent 69.98 % of the total genome (average length 878.9 bp), whereas they represent 69.93 % of the nucleotide sequence for C1, with an average gene size of 877.0 bp. For comparison, *P. penetrans,* a close relative, has 2,407 genes (average length 694 bp), representing 66.33 % of the genome (29). The genomes of both *Pasteuria* species are less dense in coding regions than related species such as *Bacillus thuringiensis* (83.28 %, (30)) or *Thermoactinomyces sp.* (86.56 %, (31)). The G+C content of both *P. ramosa* isolates is around 31 %, which is at the lower end compared to other *Bacillota spp.* (32) (*B. thuringiensis* 35 %, *Thermoactinomyces sp.* 48 %, *P. penetrans* 46 %) (29). Notably, neither a canonical CRISPR system nor prophages were identified in the two *P. ramosa* genomes using CRISPRCasFinder (33) and Phaster (34, 35) tools.

**Figure 1:**
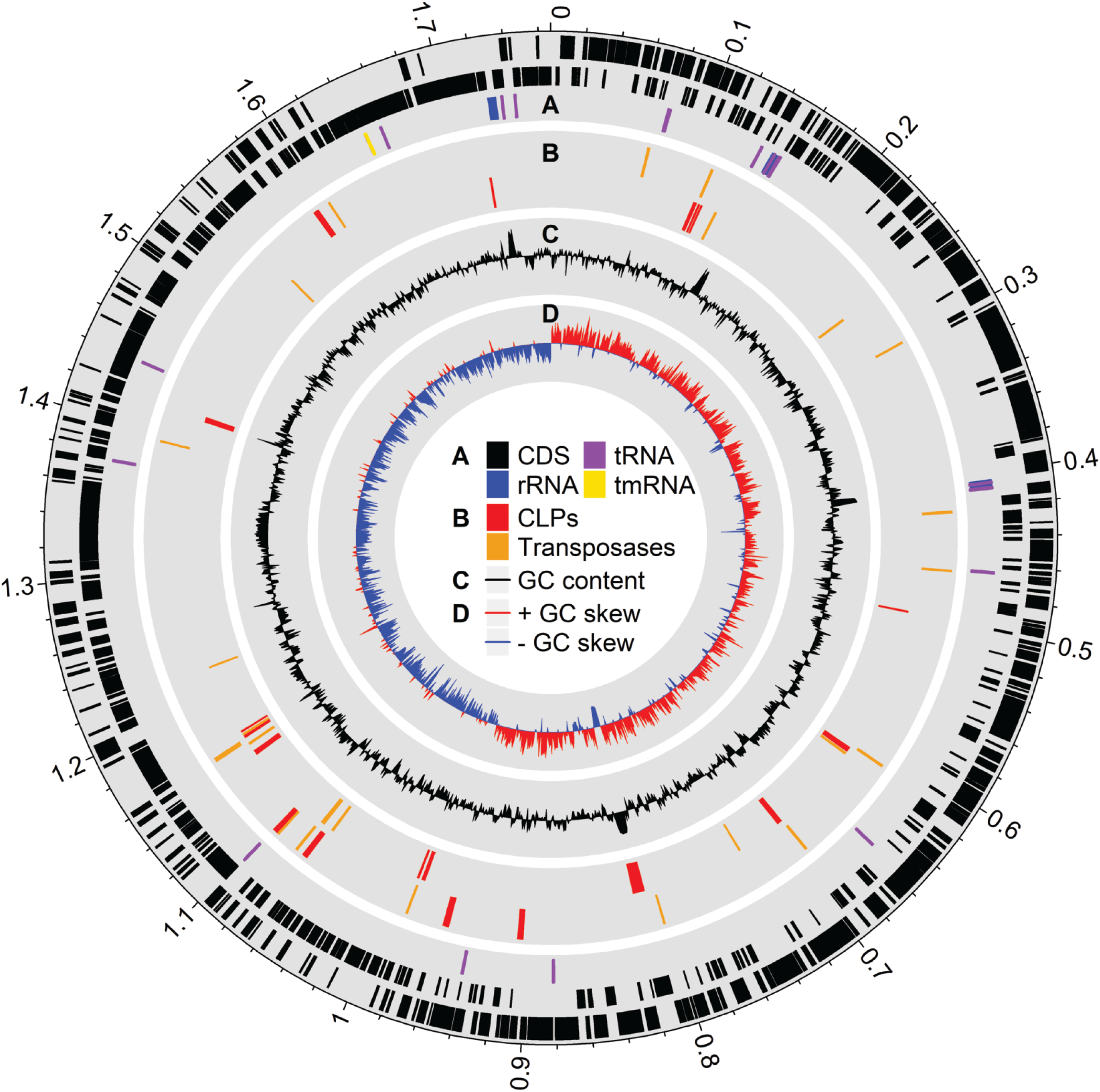
Circular map of genomic features for *P. ramosa* P21. Orientated from the origin. The total length is about 1.77 Mb (scale on the outside of the circle). (A) Structural annotation on forward (outer ring) and reverse (middle ring) strands. Structural annotation of rRNA, tmRNA and tRNA on the inner ring. (B) Location of CLP genes (red) and transposases (orange). (C) GC content of the genome scaled to the average genome-wide GC content (31.1 %). (D) GC skew. Red shading denotes GC skew values greater than 0 (i.e. overrepresentation of cytosine over guanine), whereas blue shading denotes an underrepresentation of cytosine over guanine. Graph plotted with RCircos (60).

**Table 1:** Genome metric summary of the two genomes sequenced.

|  | P21 | C1 |
| --- | --- | --- |
| Size (in bp) | 1,767,095 | 1,736,924 |
| circular (Yes/No) | Yes | Yes |
| Long read coverage | 964X | 43X |
| Short read coverage | NA | 1460X |
| GC% | 31.13 | 30.84 |
| CDS | 1407 | 1379 |
| rRNA | 8 | 8 |
| tRNA | 39 | 38 |
| tmRNA | 1 | 1 |
| BUSCO Firmicutes | 81.2 | 81.7 |
| CLPs | 39 | 43 |
| Transposases | 26 | 5 |

### Phylogenetic position and gene repertoire

A single-copy-BUSCO-gene phylogenetic tree based on the P21 genome firmly confirms the *Pasteuria* placement within the phylum *Bacillota* and the class *Bacillales* (Figure 2A, Table S1). The tree shows that the *Pasteuria* genomes diverge deeply within the larger *Bacillales*, with the closest related clade including *Thermoactinomyces sp.* (Figure 2A). *P. ramosa* and *P. penetrans* exhibit the lowest BUSCO score in the *Bacillota* phylum (81.2 % and 80.3%, firmicutes_odb10, Figure 2B), missing 38 and 39 highly conserved genes of the *Bacillota* phylum, respectively. Thirty-two of these are missing from both genomes. Fifteen of these missing genes are involved in translation, ribosomal structure, and biogenesis (Table S2). When comparing the Cluster of Orthologous Groups (COG) genome annotations between *P. ramosa, P. penetrans,* and *Thermoactinomycetes spp.,* the *Pasteuria* genomes show distinct reductions, primarily in functions related to amino acid, nucleotide, lipid metabolism, and transport (Table S3). In contrast, translation, replication machinery, and repair functions are unaffected. *P. ramosa* and *P. penetrans* exhibit a high percentage of genes (11 % and 21.8 % respectively) in species-specific orthogroups (Figure 2C). *P. ramosa* has 24 species-specific orthogroups containing a total of 159 genes; the largest orthogroup contains 30 proteins annotated as transposases (Table S4).

**Figure 2:**
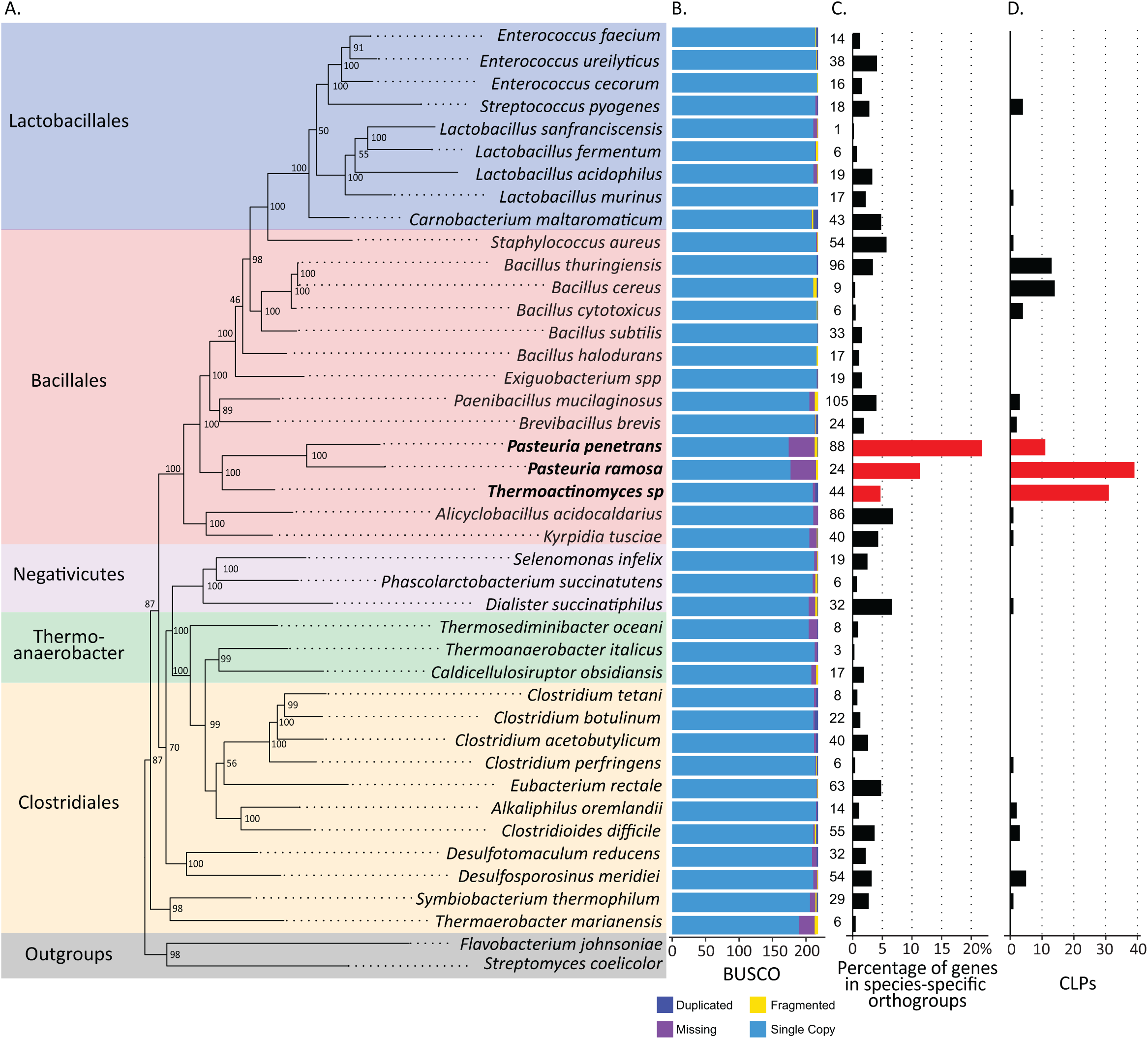
Phylogenetic tree and gene content comparison of selected Firmicutes species (based on P21 isolate’s genome). *Pasteuria ramosa* and *P. penetrans* have a high percentage of genes in species specific orthogroups as well as a high number of missing BUSCO genes. *Pasteuria ramosa* and *Thermoactinomyces sp.* have exceptionally high numbers of CLPs. Bar charts contain data for each species aligned to the corresponding species in the phylogenetic tree. (A) Phylogenetic species tree inferred using complete and single-copy BUSCO genes extracted from entire genomes of all species shown here (n = 77). Bootstrap values (percent) are indicated at the node (800 iterations). Colors indicate bacterial orders. (B) BUSCO assessment results. The reference database for BUSCO (v 5.0.0) was “firmicutes_odb10” with 218 core genes in total. Blue refers to the number of the complete single-copy orthologs; dark blue to the complete duplicated orthologs; yellow refers to fragmented or incomplete orthologs, and purple to missing orthologs. (C) Percentage of genes in species-specific orthogroups. Number to the left of the axis are the absolute numbers of genes in species specific orthogroups. (D) Number of CLPs in each species. *P. ramosa*, *P. penetrans* and *Thermoactinomyces sp.,* highlighted in red, form a monophyletic group.

### Collagen-like genes

It has been suggested that CLPs act as a bacterial adhesion, playing an important role in the attachment of *Pasteuria* spores to its host (14, 15) and making them, thus, a central mechanism in the coevolutionary process. An extensive search for CLP genes yielded a total of 48 hypothetical genes in P21. Nine of these were missing their 3’ end, making them likely fragments or pseudogenes, with one substantially longer (8 kb) than all other members of this gene family (mean = 1,082 bp, range 755 – 1949 bp). Excluding these fragments from further analysis, left us 39 putative CLP genes in P21. These CLPs contained between seven and 823 Gly-X-Y-motifs (65.36 motifs on average, an average 35.79 % of gene length). The same search in the C1 genome identified 43 CLP genes. Of these 39 (P21) and 43 (C1) CLP genes, 30 in P21 and 38 in C1 were assigned according to previous annotation, leaving nine and five genes, respectively, that had not been previously described. The two genomes had 30 CLPs in common; their identity ranged from 74.7 % to 100 %, with their position in the genome conserved (Table 2) and distributed more or less uniformly across the entire *P. ramosa* genome (Figure 1B for P21). This uniformity is also reflected in the distribution across the strands: approximately 53 % of the genes are on the leading strand. In contrast, 77 % of all genes in the *P. ramosa* genome are on the leading strand (Figure 1).

**Table 2:** *P. ramosa*-CLPs identity.

| Protein name C1 | Protein name P21 | Protein size in C1 (AA) | Protein size in P21(AA) | Percentage of identity | Clustering group |
| --- | --- | --- | --- | --- | --- |
| C1_Pcl50 | absent | 306 |  | NA | ungroup |
| C1_Pcl1a | P21_Pcl1a | 505 | 649 | 84.4 | ungroup |
| C1_Pcl02 | P21_Pcl02 | 371 | 380 | 97.6 | ungroup |
| C1_Pcl15 | P21_Pcl15 | 398 | 359 | 75.6 | 1 |
| C1_Pcl16 | P21_Pcl16 | 320 | 322 | 74.7 | 2 |
| C1_Pcl17 | P21_Pcl17 | 439 | 418 | 82.7 | 3 |
| C1_Pcl29 | P21_Pcl29 | 383 | 333 | 80.1 | ungroup |
| C1_Pcl20 | P21_Pcl20 | 253 | 253 | 100.0 | 4 |
| C1_Pcl19 | P21_Pcl19 | 254 | 254 | 99.6 | 4 |
| C1_Pcl18 | P21_Pcl18 | 302 | 302 | 99.7 | 4 |
| C1_Pcl30 | P21_Pcl30 | 251 | 251 | 99.2 | 4 |
| C1_Pcl32 | P21_Pcl32 | 256 | 256 | 100.0 | 4 |
| C1_Pcl28 | P21_Pcl28 | 412 | 412 | 98.1 | 3 |
| C1_Pcl27 | P21_Pcl27 | 315 | 315 | 100.0 | 2 |
| Fragment | P21_Pcl40 |  | 393 | NA | 1 |
| C1_Pcl03 | P21_Pcl03 | 417 | 417 | 100.0 | 3 |
| C1_Pcl04 | P21_Pcl04 | 323 | 323 | 100.0 | 2 |
| C1_Pcl05 | P21_Pcl05 | 396 | 396 | 99.7 | 1 |
| C1_Pcl33 | Absent | 193 |  | NA | ungroup |
| C1_Pcl41 | P21_Pcl41 | 1014 | 2853 | 85.6 | ungroup |
| C1_Pcl48 | Absent | 892 |  | NA | ungroup |
| C1_Pcl21 | P21_Pcl21 | 342 | 342 | 97.7 | 1 |
| C1_Pcl22 | P21_Pcl22 | 325 | 325 | 100.0 | 2 |
| C1_Pcl23 | P21_Pcl23 | 421 | 387 | 81.5 | 3 |
| C1_Pcl42 | P21_Pcl42 | 383 | 383 | 99.7 | 3 |
| C1_Pcl35 | P21_Pcl35 | 320 | 320 | 99.7 | 2 |
| C1_Pcl34 | P21_Pcl34 | 396 | 396 | 100.0 | 1 |
| C1_Pcl08 | P21_Pcl08_2 | 428 | 425 | 90.9 | 3 |
| C1_Pcl07 | P21_Pcl07_2 | 330 | 328 | 88.2 | 2 |
| C1_Pcl06 | P21_Pcl06_2 | 344 | 344 | 91.6 | 1 |
| C1_Pcl36 | Absent | 335 |  | NA | 1 |
| C1_Pcl37 | Absent | 326 |  | NA | 2 |
| C1_Pcl38 | Absent | 421 |  | NA | 3 |
| C1_Pcl14 | Absent | 345 |  | NA | 1 |
| C1_Pcl13 | Absent | 323 |  | NA | 2 |
| C1_Pcl12 | Absent | 421 |  | NA | 3 |
| Absent | P21_Pcl45 |  | 341 | NA | 1 |
| Absent | P21_Pcl46 |  | 323 | NA | 2 |
| Absent | P21_Pcl47 |  | 422 | NA | 3 |
| C1_Pcl11 | Absent | 335 |  | NA | 1 |
| C1_Pcl10 | Absent | 331 |  | NA | 2 |
| C1_Pcl09 | Absent | 420 |  | NA | 3 |
| C1_Pcl26 | P21_Pcl26 | 412 | 411 | 92.0 | 3 |
| C1_Pcl25 | P21_Pcl25 | 316 | 317 | 83.0 | 2 |
| C1_Pcl24 | P21_Pcl24 | 378 | 333 | 84.2 | 1 |
| C1_Pcl49 | P21_Pcl43 | 1168 | 539 | 71.6 | ungroup |
| C1_Pcl31 | P21_Pcl31 | 231 | 255 | 88.2 | ungroup |
| Absent | P21_Pcl44 |  | 558 | NA | ungroup |

Many CLP genes are organized in triplets, usually with one gene from each of three groups (referred to as group 1, 2, and 3 (Figure 4, 35) clustering tightly together, all in the same order and orientation (Figure 3). Among the eight newly detected CLP genes in P21, three form a triplet (*Pcl45-46-47*), two complete triplets that were previously identified as incomplete (*Pcl40*, *Pcl42*), while the others remain independent of any triplet or clustering group (*Pcl43*, *Pcl44*). The new CLP genes detected in C1 are all ungrouped except *Pcl42*, which completes a previously recorded triplet.

**Figure 3:**
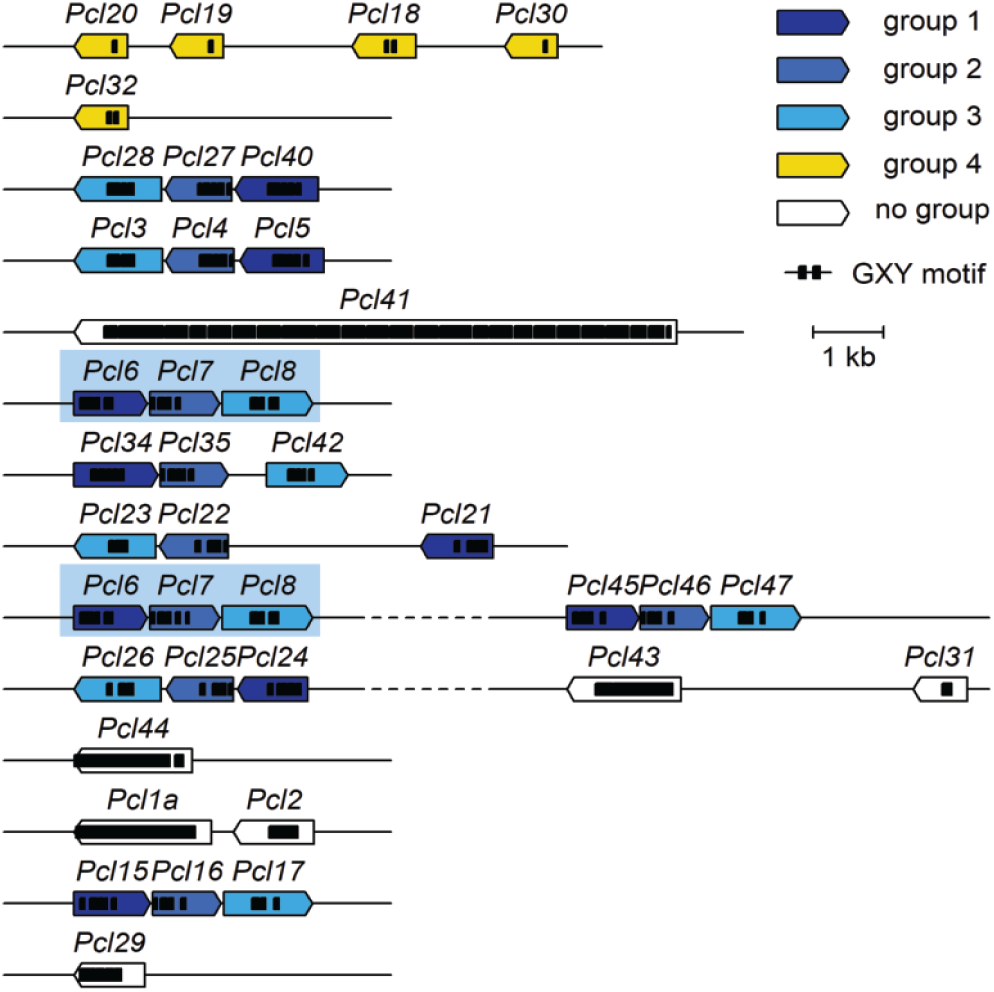
Organization of *P. ramosa*-CLP genes within the genome. Fragments of CLP genes are not plotted. Colours indicate groupings of genes from the gene tree in Fig. 4. Triplets are formed by 3 CLP genes and have a consistent structure - composed of one gene from each group 1, 2 and 3 in the same order and orientation.

**Figure 4:**
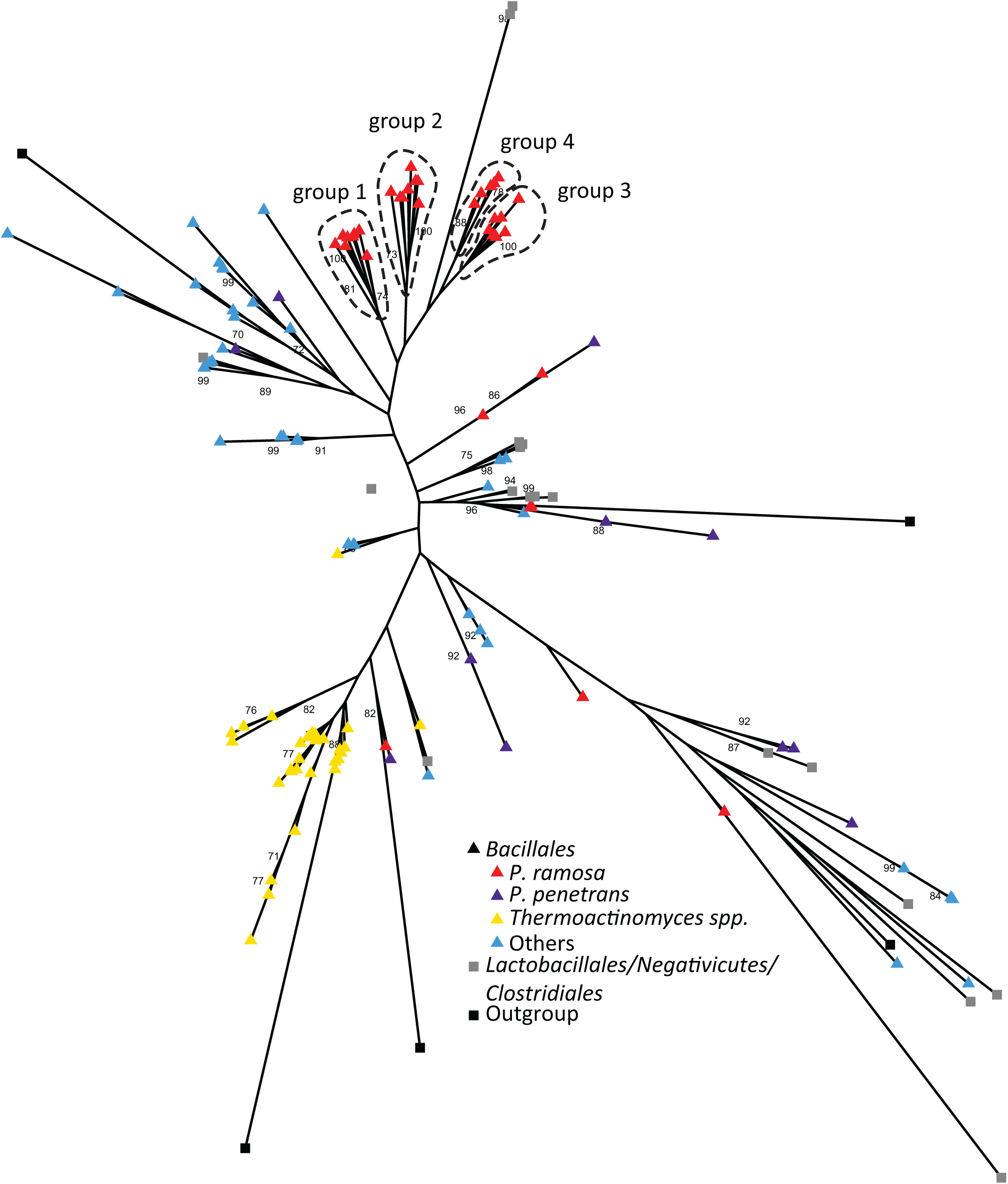
Firmicutes CLP gene tree. Maximum-likelihood tree based on CLP genes sequences in the Firmicutes species included in Fig. 2. While *P. ramosa*-CLP genes (red) and *Thermoactinomyces sp.*-CLP genes (yellow) are strongly clustered, the CLP genes of the other species (grey) including *P. penetrans* (purple) do not show a clear clustering pattern. Four clusters of *P. ramosa* CLP genes (group 1-4), correspond to the four groups shown in Fig. 3. Bacillales order is indicated by triangles; the other orders, by squares. Bootstrap support values over 70 are displayed on the nodes.

One CLP-triplet (*Pcl45-46-47*) detected in the *P. ramosa* P21 isolate exhibits a high homology (70.33 % to 89.81 %) with two other triplets (*Pcl12-13-14* and *Pcl36-37-38*) found in the C1 isolate, suggesting that copy/paste mechanisms may be involved in the evolutionary process for these triplets. Triplet *Pcl6-7-8* is also duplicated, with genes *Pcl6* and *Pcl7* being perfectly identical, and *Pcl8* having one synonymous mutation. This triplet’s duplication is part of the duplication of a 6.5 kb stretch of DNA (confirmed by sequencing (Figure 3, File S1 and S2)). One triplet, *Pcl21-22-23*, deviates from the triplet rule in that it includes two genes between the *Pcl21* and *Pcl22*: one coding for a full-length IS5 transposase and the other for a ferritin-like domain-containing protein.

Within the *Bacillota* phylum, CLPs are found predominantly in the *Bacillales* order (13, 36) with large numbers of them located mainly, though not exclusively, in the *Pasteuria/Thermoactinomyces* clade (as Figure 2D illustrates). To determine if these CLP genes represent an ancient radiation of this gene family or if they radiated within species, we constructed a maximum likelihood tree of all CLP genes identified in the *Bacillota* (Figure 4). The *P. ramosa* CLP genes primarily form a tight cluster (indicated in red in Figure 4), displaying a clear sub-structure consisting of four groups. The pattern correlates well with the clustering pattern detected by McElroy et al. (2011)(26). However, some *P. ramosa* CLP genes are scattered throughout the entire tree (Figure 4), suggesting they may have diversified much earlier than the CLP genes of groups 1 to 4. The CLP genes of *P. penetrans* are also scattered across the entire tree without recognizable clustering. Interestingly, we identified 47 hypothetical CLP genes in *Thermoactinomyces sp*. that cluster almost perfectly together (with one exception), indicating species-specific radiation.

### Genome plasticity and synteny of the two isolates

The largest orthogroup specific to *P. ramosa* is annotated as containing transposases. Transposases play a role in genome rearrangement by promoting inversions, duplications, deletions and transpositions. These changes can affect significant portions of genomes and may alter the infectivity or pathogenicity of pathogenic bacteria (37).

At first, 86 and 54 hypothetical transposases were identified in the P21 and C1 genomes. However, manual curation reduced this to 26 and five transposases, respectively. These transposases fall into two families (IS701 and IS5, Table 3): for P21, nine IS701 members share pairwise identities ranging from 98.35 % to 100 %, with two pairs being perfectly identical (Figure S3A). Similarly, ten IS5 proteins exhibit pairwise identities ranging from 97.6 % to 100 %, with two and five perfectly identical (Figure S3B). This pattern suggests recent duplication or burst events. In contrast, none of the complete transposases in the C1 genome are identical. Strikingly, the pairwise identities in the IS701 family between the proteins of each isolate are lower (from 84.18 % to 89.81 %), except for one pair where the identity reaches 99.05%.

**Table 3:** Transposases.

| Transposase family | Number in P21 | Number in C1 | Identical genes in P21 | Identical genes in C1 |
| --- | --- | --- | --- | --- |
| IS5 | 13 | 2 | [NEKOFPGG_00195, NEKOFPGG_00241, NEKOFPGG_00993, NEKOFPGG_01050, NEKOFPGG_01259] | NA |
|  |  |  | [NEKOFPGG_01316 and NEKOFPGG_01368] | NA |
| IS701 | 13 | 3 | [NEKOFPGG_00483 and NEKOFPGG_00896] | NA |
|  |  |  | [NEKOFPGG_00494 and NEKOFPGG_00902] | NA |

To investigate the conservation of genome structure between the two isolates, we constructed a dot-plot of the C1 genome against the P21 genome. This analysis revealed a large structural variant between them: a genome fragment, 174 kb in size, is inverted in one of the isolates but otherwise identical (Figure 5A). This inversion contains 3 CLP triplets (*Pcl6-7-8*, *Pcl34-35-42*, *Pcl23-22-21*). Additionally, a further structural difference was found in a region of approximately 40 kb (Figure 5B) that in the P21 isolate contains two CLP triplets (the second copy of the duplicated *Pcl6-7-8* triplet and *Pcl45-46-47*), but in C1 contains three CLP triplets (*Pcl36-37-38*, *Pcl14-13-12*, and *Pcl11-10-9*). Within this region, the P21 genome contains six complete transposases, whereas the C1 genome contains only two, along with one fragment. Moreover, some structural variations between P21 and C1 localize within a region of 100 kb (P21 1.592 to 1.622 Mb, C1: 1.562 to 1.592 Mb) (Figure S2). In this region, the P21 isolate contains one full IS5 transposase and eight fragments of transposases, while the C1 genome contains two full IS701 transposases and three fragments of other transposases, within the same region a 6 kb inversion was found.

**Figure 5:**
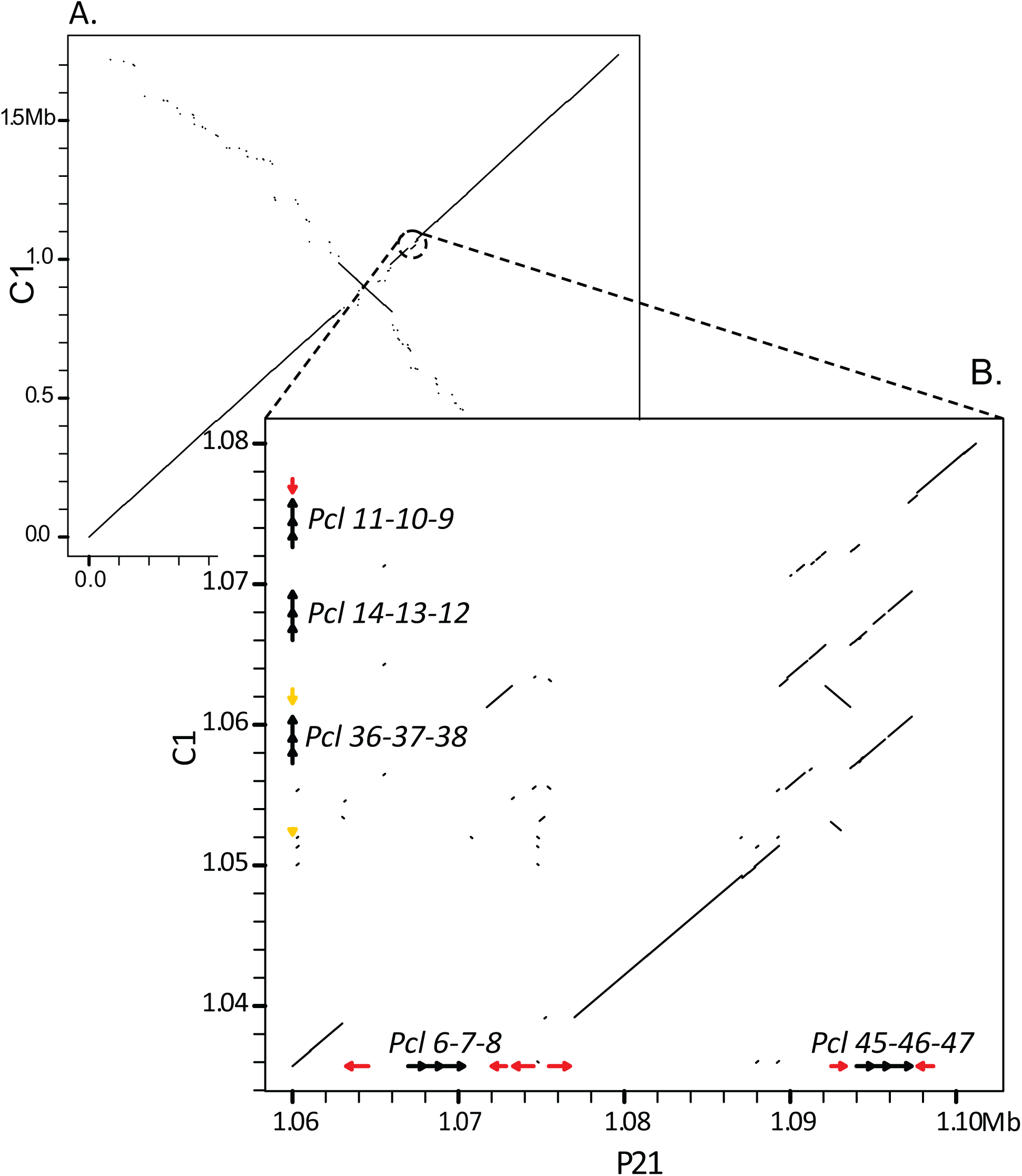
Comparison of genomes for *Pasteuria ramosa* isolates C1 and P21 using a dot plot. The two genomes are very similar, except for one genome rearrangement and one variable region. (A) Dot plot alignment of P21 isolate (horizontal axis) versus C1 isolate (vertical axis). An inversion of approximatively 150 kb is visible, as well as a more variable region (encircled) slightly upstream of it. (B) Close-up of the previous dot plot (P21 [1.06..1.105 Mb] and C1 [1.035 .. 1.08 Mb]). CLPs positions are indicated by black arrows; transposases locations are indicated by red arrows for the complete genes and in yellow for the fragments. *Pcl45-46-47* in P21 has a high homology with *Pcl36-37-38* and *Pcl14-13-12* in C1. The triplet *Pcl6-7-8* in P21 does not exist in C1, and triplet *Pcl11-10-9* does not exist in the zoomed-in region of P21. Note: a nearly perfect copy of triplet *Pcl67-8*, exists at another place in both genomes.

## Discussion

Our analysis allowed us to place the genus *Pasteuria* firmly within the *Bacillales* clade, where it forms a monophyletic group together with the free-living *Thermoactinomyces*, with which it shares unusual growth characteristics. Both genera produce branching hyphae-like structures during growth (38, 39). The parasitic lifestyle of *Pasteuria* has drastically impacted its genomes, reducing amino acid, nucleotide, lipid metabolism and transport capabilities. It is likely they may obtain these compounds directly from their hosts. The sister species of *P. ramosa, P. penetrans,* is also divergent, with 11 % and 21.8 % of its genes present in species-specific orthogroups. *P. ramosa* and *P. penetrans* parasitize a crustacean and a nematode, respectively, and have consequently evolved specific functions or genes tailored to their individual hosts.

### CLP coevolution and diversification

*P. ramosa* and *D. magna* share an ancient history of coevolution (27, 40). Multiple loci driving resistance to attachment have been detected in the host (22, 23) that appear to be evolving in an antagonistic, coevolutionary process consistent with the Red Queen model (41). The *P. ramosa* CLP family of proteins is suspected to be responsible for attachment to the host cuticle (27, 42). Phylogenetic analysis of this family of proteins in *Pasteuria* and related species has revealed that some CLPs radiated extensively in *P. ramosa* and *Thermoactinomyces*. At the same time, this pattern is absent in most other species, including *P. penetrans* (Figure 4). In *P. ramosa*, most CLP genes cluster in four tight groups, each radiating from a single ancestor. Additionally, most CLP genes in *P. ramosa* are organized in triplets, comprising members from groups 1, 2, and 3 - three of the four tight clusters - in the same order and orientation. This pattern suggests they may have arisen from duplication events of entire triplets. We speculate that the radiation and triplet organization of CLP genes in *P. ramosa* is related to their attachment function and that the parasite needs to diversify its repertoire of proteins to facilitate attachment and infection of the host. Consistent with this, *Pcl7*, the only CLP gene whose function is currently characterized in *P. ramosa*, is part of a triplet.

However, some *P. ramosa* CLP genes do not conform to this triplet organization and display a distinct evolutionary pattern (Figure 4). Among them, some show a tight clustering, such as group 4 (Figure 4), and high conservation, suggesting a potential structural and essential role. Conversely, some *Pasteuria* CLPs are widely distributed across the CLP gene tree, indicating they may not be subject to strong selection.

The two isolates sequenced here, P21 and C1, differ in their infectivity to a panel of host genotypes, as well as in the CLPs family members they contain. While we found more than 50 CLP family members, the genomes of the two isolates contain only 39 and 43 of them, respectively. These two sets of CLPs are largely overlapping, with 30 CLPs in common (Table 2). Thus, the composition seems variable, as might be expected if CLPs are subject to a selective process associated with coevolution. More precisely, the diversification of CLP triplets in *P. ramosa* seems to be ongoing, as evidenced by a recent duplication of CLP triplet *Pcl6-7-8* in P21 and the divergence of others (*Pcl12-13-14* and *Pcl36-37-38* in C1, and *Pcl45-46-47* in P21). A mechanism that allows for diversification and radiation of a gene family is gene duplication, followed by mutation and selection. We discovered 26 and five transposases in the two studied *P. ramosa* genomes, primarily within two specific families (IS5 and IS701). Generally, a genome with an burst of transposases is consistent with other bacterial systems that have recently undergone a host-restricted lifestyle (43). However, since *P. ramosa* and *D. magna* have been coevolving for at least a million years (27), these transposases might have been conserved for another purpose. Here, we speculate that they may contribute to the continuous diversification of the CLPs family, a pattern that is consistent with the variation in the number of transposases from one isolate to another. Such a pattern could result, presumably, in a constant birth and death process of genes, leaving behind a graveyard of non-functional genes or gene fragments. Indeed, in C1 we find many fragments and pseudogenes of transposases: out of 20 putative IS701 genes, 10 are only fragments, and 10 are non-functional proteins due to mutations. Importantly, we do not focus on the pseudogenisation of other gene families in the present study. This high activity of transposases might also explain the unexpectedly low gene density of this parasite’s genome.

On the other hand, the high number of transposases might lead one to expect more structural variations between the two isolates, which we did not see. This result could simply be the result of comparing only two genomes; studying additional genomes may reveal more structural variation and less conservation of genome structure.

## Conclusion

In summary, our study highlights the dynamic interplay of gene duplication, mutation, and selection within the CLP family of *Pasteuria ramosa*, contributing to its genotypic specificity in host attachment. The observed transposase activity, while potentially linked to gene diversification, poses intriguing questions about its role in the long-term coevolutionary history with *Daphnia magna*. Future investigations with a broader genome sampling may shed further light on structural variations and genomic adaptations in this intricate host-parasite system, ultimately advancing our understanding of the coevolutionary arms race between *P. ramosa* and its host.

## Material and methods

### Pasteuria ramosa system

*Pasteuria ramosa* (Bacteria: Bacillota) is an endospore-forming obligate parasite of the planktonic crustacean *Daphnia* first observed by Metchnikoff (44). If the host and the pathogen genotype are compatible, attachment is possible. An attached bacterium penetrates the cuticle and enters the host’s body cavity where it replicates, eventually producing millions of spores that are released after the death of the host. Infection results in host castration and a shortened lifespan (21). Transmission is exclusively horizontal. Although *P. ramosa* is so far unculturable in vitro, it can be passaged repeatedly through susceptible hosts, and, as a result of the distinct bottleneck that takes place as part of the transmission process, pure lineages of single genotypes can be generated.

High-quality DNA was produced by focusing on the immature stage of the spores, which do not yet have an exosporium and can be lysed without causing DNA damage. However, immature spores cannot be separated from the host tissue easily. Here, we used exclusively immature spores from infected hosts to obtain DNA. We focus on two *P. ramosa* isolates commonly used in the lab, P21 and C1. P21 was first isolated from an infected *D. magna* individual collected in a pond in Switzerland (Aegelsee, Switzerland, GPS: 47.558, 8.863), and C1 from an individual collected in a pond at the Moscow Zoo (Moscow, Russia, GPS: 55.763, 37.582)(19).

### Infection trials and collection of infected *D. magna*

*Daphnia magna* clones were kept under standardized conditions in artificial culture medium (ADaM, (45)) at 20 °C. Infection trials were carried out according to a previously published protocol (46). The isolate C1 was propagated in the host clone HU-HO-2 (Bogarzo-to, Hungary, GPS: 46.8, 19.1333), whereas the P21 isolate was propagated in the host clone t0_9.3_7 (Aegelsee). At 14 days post-infection, when we reached the grape-seed stage (the "immature spores") (21), we began collecting infected animals. Twelve hours before collection, infected *D. magna* were individually placed into 100-mL jars with 50 mL of freshly filtered ADaM (20-µm filter). *D. magna* microbiota, undigested food, and the micro-biofilm covering the carapace of the animal are significant sources of DNA contamination. To reduce gut DNA contamination, each animal was fed dextran beads (Sephadex G-25 by Sigma Aldrich, 5g/L) every six hours, a treatment that clears the gut. We also observed individuals every hour to see if they had molted, which happens every three to four days when *D. magna* individuals shed and produce a new carapace not yet covered by a bacterial biofilm. Molted individuals were collected and pooled in a 100-mL jar containing freshly filtered ADaM, and stored at 4 °C. Batches of approximately 20 individuals were transferred to a 1.5-mL microcentrifuge tube with 100 µL of G2 Buffer (lysis buffer from Genomic DNA Buffer Set, Qiagen ID: 19060) and carefully crushed with a plastic pestle less than four hours after their collection.

### DNA extraction and sequencing

DNA was extracted from 91 P21- and 88 C1-infected animals using a Qiagen Genomic Tips kit (Qiagen ID: 10223). The P21 DNA sample was sent to Genomic Facility of ETH D-BSSE (Basel, Switzerland) for sequencing where a PacBio SMRTbell library was prepared following DNA size selection with the BluePippin (DNA > 8 kb). The resulting sample was run on a single PacBio Sequel I SMRTcell. In total, ∼ 5.1 Gbp of continuous long-read (CLR) sequence data (∼2.4 million reads) were obtained, with mean subread length of 7.3 kb (N50 = 9.6 kb).

The C1 genotype was sequenced using the Oxford Nanopore MinION technology. The library for MinION was constructed using a ligation kit (SQK-LSK110, Oxford Nanopore Technology, Oxford, UK) and then analyzed using two FLOMIN106 flow cells (v9.4.1). In total, 8,732,179 reads were collected for a read length average of 562bp. The raw FAST5 data were basecalled using Guppy v6.0.1 with the super accurate mode. We also used high-accuracy Illumina data, generating 17,875,920 Illumina MiSeq PE-250 bp reads based upon a Nextera XT DNA preparation kit. Sequencing was performed at the Department of Biosystems Science and Engineering, ETH-Zurich in Basel, Switzerland.

### Genome assemblies and annotations

#### P21 isolate assembly

We initially mapped all the reads to a draft genome of *P. ramosa* C1 based only on Illumina data (unpublished) and filtered out all the unmapped reads (236,507 mapped reads). We then used the Flye assembler (v2.8.3-b1695, 49) with PacBio reads specific parameters to construct the primary assembly. To correct errors in the primary assembly, we used the Arrow pipeline from Genomic consensus toolkit to polish the genome (https://github.com/PacificBiosciences/GenomicConsensus). The assembled genome is 1.77 Mb and circular. To evaluate the relative biological completeness of this new genome draft, we used BUSCO v5.0.0 (48) and the lineage database Firmicutes_odb10 (version 2021-02-23).

For the annotation, including gene prediction, tRNA and rRNA detection, we used Prokka v1.14.6 (49) with the whole bacterial non-redundant protein database from NCBI (RefSeq Release 204, 14^th^ Jan. 2021). We paid special attention to the annotation of the *Pcl* genes. We downloaded the 38 known CLP genes sequences from the UniProt website and used this file as a reference set of trusted proteins to annotate with Prokka and the option “`--protein`”. In addition, Prokka gene predictions were examined with HMMER 3.3.2 (http://hmmer.org/, (50)) and a Pfam collagen domain (PF01391) to search for additional potential CLPs. The list of CLPs was manually curated, and fragments were removed. The origin of the chromosome was determined with the web tool Ori-finder (March 2021, 53). The replication origin was localized at the position at the base 444,423bp.

#### C1 isolate assembly

We initially mapped the short and the long reads to a draft genome of *P. ramosa* C1 as described above (unpublished) and filtered out all unmapped reads. Then the ONT long-read dataset was subset according to the read length, with only reads > 5 kb kept for assembly (14,420 reads for an average coverage of 42.6 X). For the short reads, we used only the reads that mapped to the draft genome (9,761,270 PE reads and 548,741 SE reads). A hybrid assembly was made using Unicycler v0.5.0, (52), resulting in a circular chromosome of 1.72 Mb. The origin of the chromosome was determined with the web tool Ori-finder (51), setting the origin at base 546,143 bp. The annotation pipeline used for P21 was also applied to C1.

#### Transposase annotations

Transposases were annotated using the webtool ISsaga (http://issaga.biotoul.fr/, 54). The list of putative transposases was manually curated: to be considered as complete, a transposase should be composed of a DDE domain, annotated as IS5 or IS701, and be longer than 250 AA for IS5 and 300 AA for IS701. Candidates with similarities to accessory genes, but without associated transposases, were excluded because we do not consider them as transposase. The proteins were aligned, and the pairwise identity was calculated using Clustal Omega (webtool 2022, 56).

### Phylogenetic tree and comparative genomics

To assess the phylogenetic relationship between the *Pasteuria* genus and the *Bacillota* phylum, we downloaded complete genomes and proteomes from a wide and representative sampling of *Bacillota* (Table S1), including a genome draft of the congeneric *P. penetrans* (29). As only the genome of *P. penetrans* was available, we annotated it using the same Prokka pipeline we used for *P. ramosa*. Other genomes used here are a subset of the data set used by (55). BUSCO and the lineage Firmicutes_odb10 (version 2021-02-23) were used to assess genome completeness and to extract all 77 single-copy BUSCO genes present in all species. These were then aligned separately using Prank v.170427 and concatenated with “catfasta2phyml.pl” script (https://github.com/nylander/catfasta2phyml). PartitionFinder was run to determine the best partition (56, 57), followed by RAxML-NG v1.1, which was run according to this partition with the `--bs-trees autoMRE` option (57). The tree converged after 800 iterations. Proteomes were clustered using OrthoFinder (2.5.2) and functionally annotated using InterProScan (v5.52-86.0). COG annotation was carried out for each genome using eggNOG-mapper (webtool 2022, 58).

### CLPs maximum likelihood tree

In order to detect Collagen Like Proteins (CLPs), a profile-based homology search of GXY motifs was performed using the HMMER package with PF01391 motif (http://hmmer.org/, (50). Only proteins longer than 150 AA and starting with a methionine were included. The 143 resulting CLP candidates were aligned with Prank v.170427, and Maximum Likelihood tree was performed with RAxML-NG with the model PMB+IO+FO. The tree converged after 650 iterations.

### Genome-genome alignment

The full genome-genome alignment of P21 and C1 was done with the tool Lastz_32 (59) according to the following options `--notransition --step=20 --gapped --chain`. The zoom-in the region P21 [1060000..1105000] and C1[1035000..1080000] was carried out following the options `--notransition -- step=20 --nogapped`. Graphs were plotted with base R 4.1.0 (R Core Team, 2021).

## Data Availability Statement

Raw data is deposited at the NCBI SRA database, while the assembled genome as well as the predicted set of protein sequences are available at NCBI GenBank (PRJNA1064691 and PRJNA1064693) and at https://figshare.com/s/8900ec07c57f352cacbf. Scripts are available at https://github.com/AThivolle/Thivolle_et_al_2024/. They will be publicly available upon acceptance.

## Author Contributions

All authors designed the study. AT and MP performed the molecular work. AT and PDF analysed the data. AT, ED, and PDF wrote the manuscript. All authors reviewed the manuscript.

## Acknowledgements

We thank Jürgen Hottinger, Urs Stiefel and Michelle Krebs for help in the laboratory. We thank members of the Ebert group for providing feedback on the study and the manuscript.

## Supplementary

### Figures

**Figure S1: Circular map of complete *P. ramosa* C1 genome.** Orientated from the origin. Total length 1.73 Mb (scale on the outside of the circle). (A) Structural annotation on forward (outer ring) and reverse (middle ring) strands. Structural annotation of rRNA, tmRNA and tRNA on the inner ring. (B) Location of CLP genes and transposases. (C) GC content of the genome scaled to the average genome-wide GC content (31.1 %). (D) GC skew. Red shading denotes GC skew values greater than 0, overrepresentation of cytosine over guanine; blue shading denotes underrepresentation of cytosine over guanine. Graph plotted with RCircos (60).

**Figure S2: C1 versus P21 variable region containing a high number of transposase fragments.** Dot plot alignment of P21 isolate (horizontal axis) versus C1 isolate (vertical axis) zoomed in on the region [1.59 .. 1.624 Mb] for P21 and [1.56 .. 1.594 Mb] for C1. This region lack of synteny between the two isolates and contains 10 fragments and one complete transposase in P21, compared to 3 fragments and 2 complete transposases in C1. There is also the inversion of a 6 kb fragment.

**Figure S3: Pairwise identity between the two main transposases family IS701 and IS5.** P21 genes are coded NEKOFPGG_XXXXX, while C1 genes are coded AKNPHEEL_XXXXX (with a grey background). A. IS701 family. Nine P21 proteins have a high similarity (98.4 to 100 %) while between isolates, the identity is pretty low (84.2 to 89.8 %) except for one couple. B. IS5 family. According to the pairwise identity and the alignment, the IS5 family can be subdivided in two subfamilies as presented in the figure. The proteins are more conserved between isolates in this family.

### Tables

**Table S1:** Accession number of the species genome and proteomes used in the study.

| Specie Name | Proteome ID | Genbank Accession Number |
| --- | --- | --- |
| *Alicyclobacillus acidocaldarius* | UP000001917 | GCA_000024285.1 |
| *Alkaliphilus oremlandii* | UP000000269 | GCA_000018325.1 |
| *Bacillus cereus* | UP000001417 | GCA_000007825.1 |
| *Bacillus cytotoxicus* | UP000002300 | GCA_000017425.1 |
| *Bacillus halodurans* | UP000001258 | GCA_000011145.1 |
| *Bacillus subtilis* | UP000001570 | GCA_000009045.1 |
| *Bacillus thuringiensis* | UP000011719 | GCA_000341665.1 |
| *Brevibacillus brevis* | UP000001877 | GCA_000010165.1 |
| *Caldicellulosiruptor obsidiansis* | UP000000347 | GCA_000145215.1 |
| *Carnobacterium maltaromaticum* | UP000000212 | GCA_000317975.2 |
| *Clostridioides difficile* | UP000001978 | GCA_000009205.2 |
| *Clostridium acetobutylicum* | UP000000814 | GCA_000008765.1 |
| *Clostridium botulinum* | UP000001986 | GCA_000063585.1 |
| *Clostridium perfringens* | UP000000818 | GCA_000009685.1 |
| *Clostridium tetani* | UP000001412 | GCA_000007625.1 |
| *Desulfosporosinus meridiei* | UP000005262 | GCA_000231385.3 |
| *Desulfotomaculum reducens* | UP000001556 | GCA_000016165.1 |
| *Dialister succinatiphilus* | UP000003277 | GCA_000242435.1 |
| *Enterococcus cecorum* | UP000017415 | GCA_000492155.1 |
| *Enterococcus faecium* | UP000325664 | GCA_008728475.1 |
| *Enterococcus ureilyticus* | UP000094469 | GCA_001730315.1 |
| *Eubacterium rectale* | UP000001477 | GCA_000020605.1 |
| *Exiguobacterium spp.* | UP000000716 | GCA_000023045.1 |
| *Flavobacterium johnsoniae* | UP000006694 | GCA_000016645.1 |
| *Kyrpidia tusciae* | UP000002368 | GCA_000092905.1 |
| *Lactobacillus acidophilus* | UP000006381 | GCA_000011985.1 |
| *Lactobacillus fermentum* | UP000094714 | GCA_001742205.1 |
| *Lactobacillus murinus* | UP000051612 | GCA_001436015.1 |
| *Lactobacillus sanfranciscensis* | UP000001285 | GCA_000225325.1 |
| *Paenibacillus mucilaginosus* | UP000007523 | GCA_000250655.1 |
| *Pasteuria penetrans* | NA | GCA_900538055.3 |
| *Pasteuria ramosa* | NA | NA |
| *Phascolarctobacterium succinatutens* | UP000004923 | GCA_000188175.1 |
| *Selenomonas infelix* | UP000004129 | GCA_000234095.1 |
| *Staphylococcus aureus* | UP000008816 | GCA_000013425.1 |
| *Streptococcus pyogenes* | UP000000750 | GCA_000006785.2 |
| *Streptomyces coelicolor* | UP000001973 | GCA_000203835.1 |
| *Symbiobacterium thermophilum* | UP000000417 | GCA_000009905.1 |
| *Thermaerobacter marianensis* | UP000008915 | GCA_000184705.1 |
| *Thermoactinomyces sp* | UP000183152 | GCA_900119485.1 |
| *Thermoanaerobacter italicus* | UP000001552 | GCA_000025645.1 |
| *Thermosediminibacter oceani* | UP000000272 | GCA_000144645.1 |

**Table S2:**
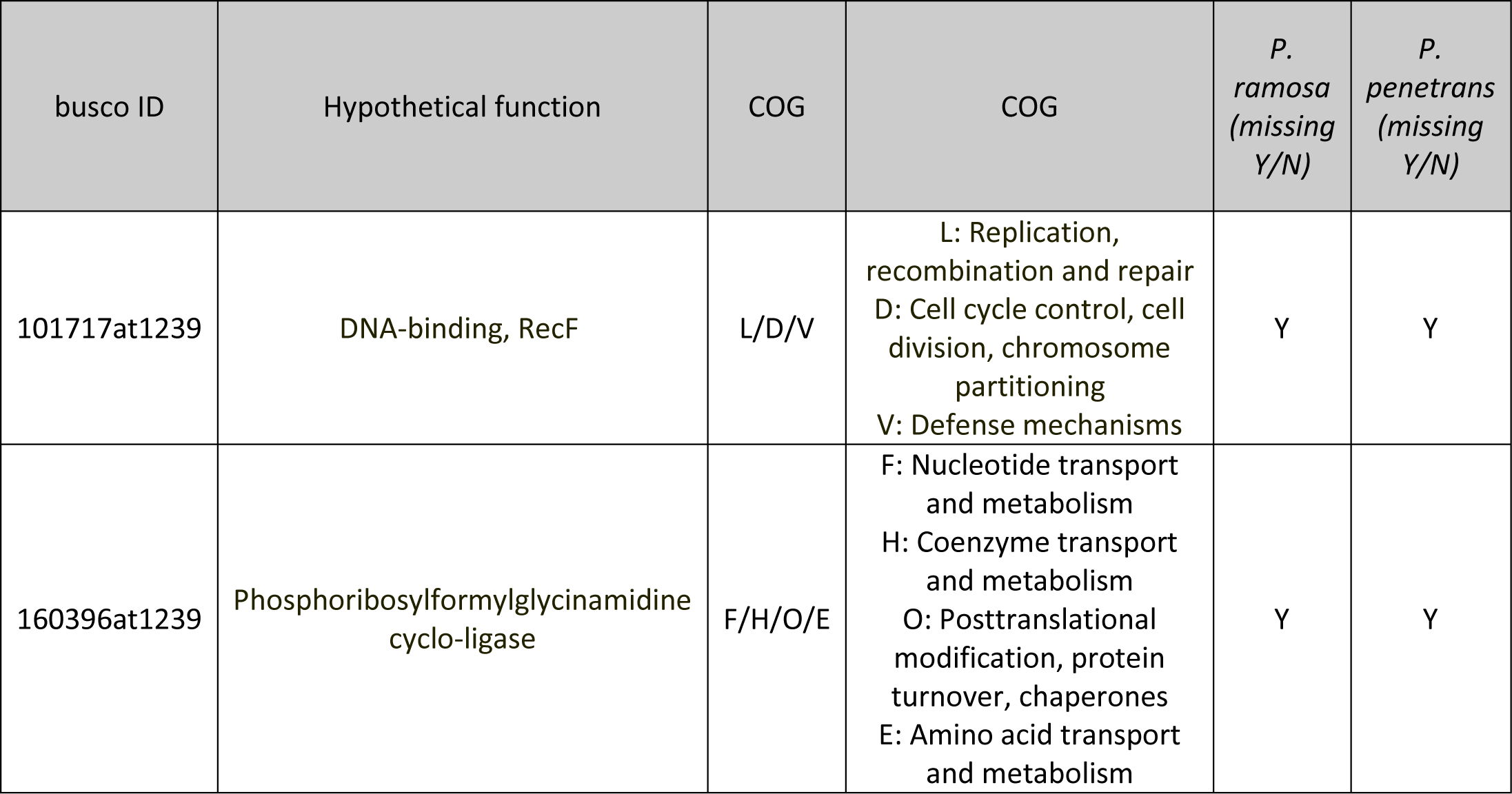
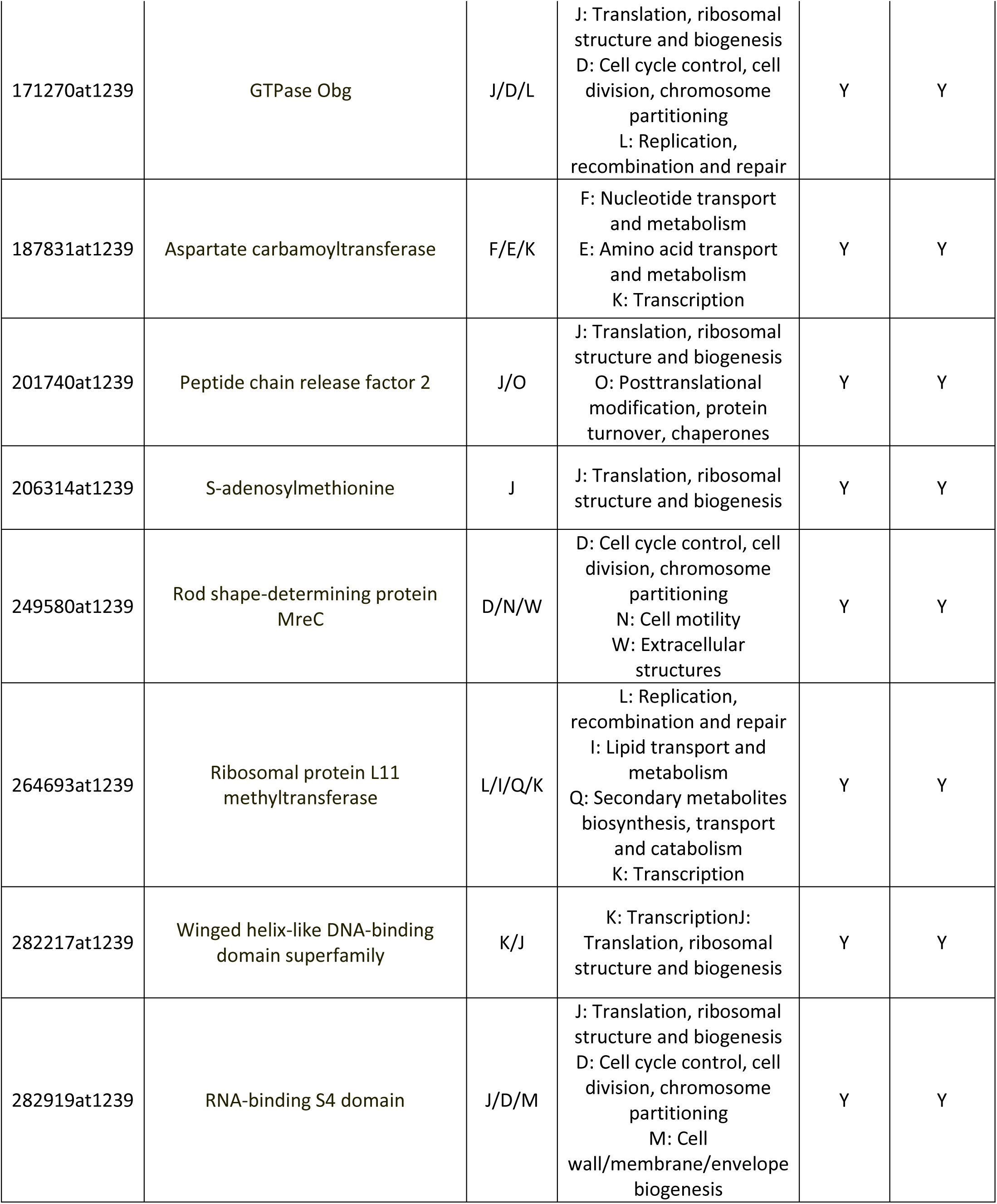
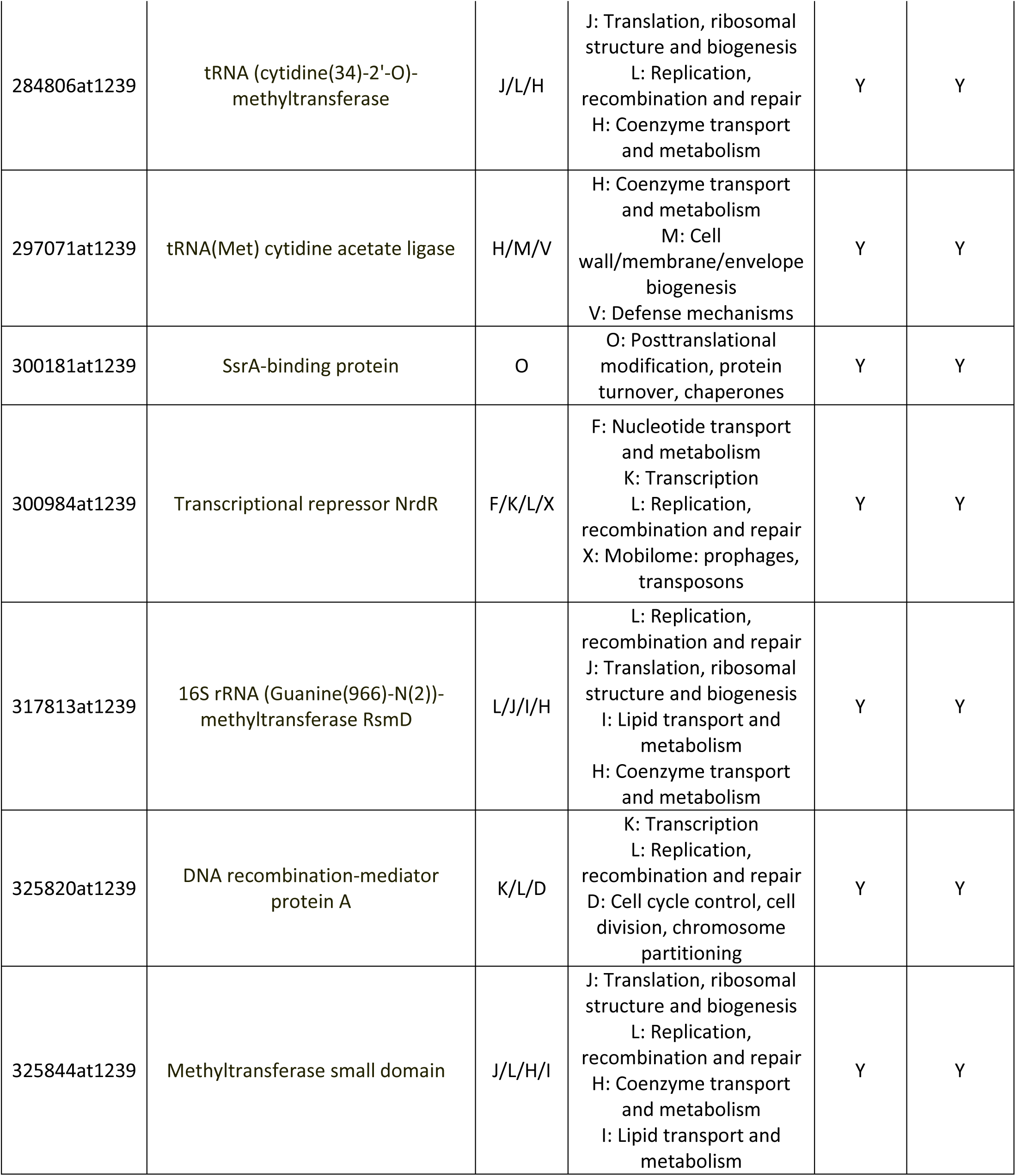
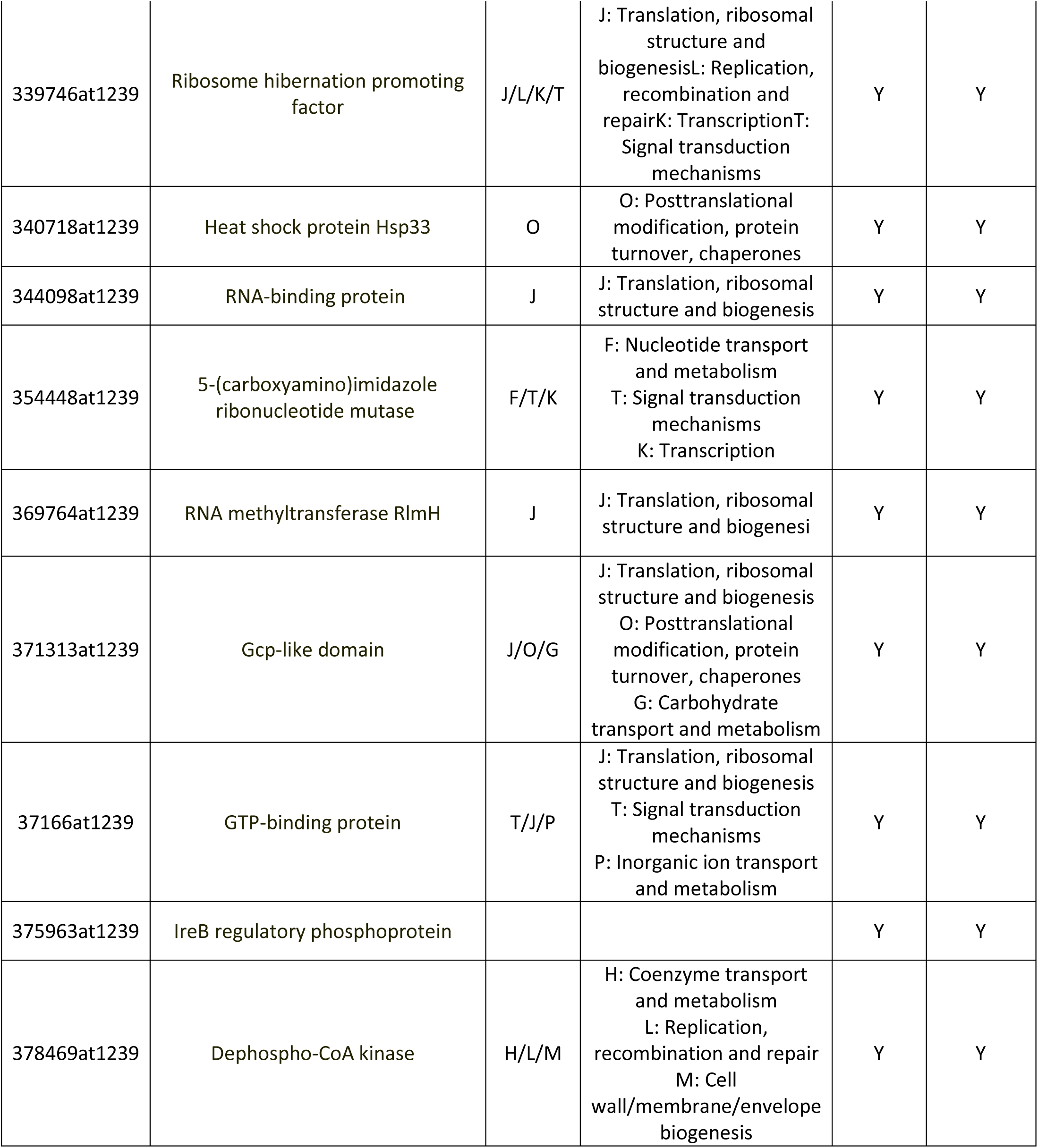
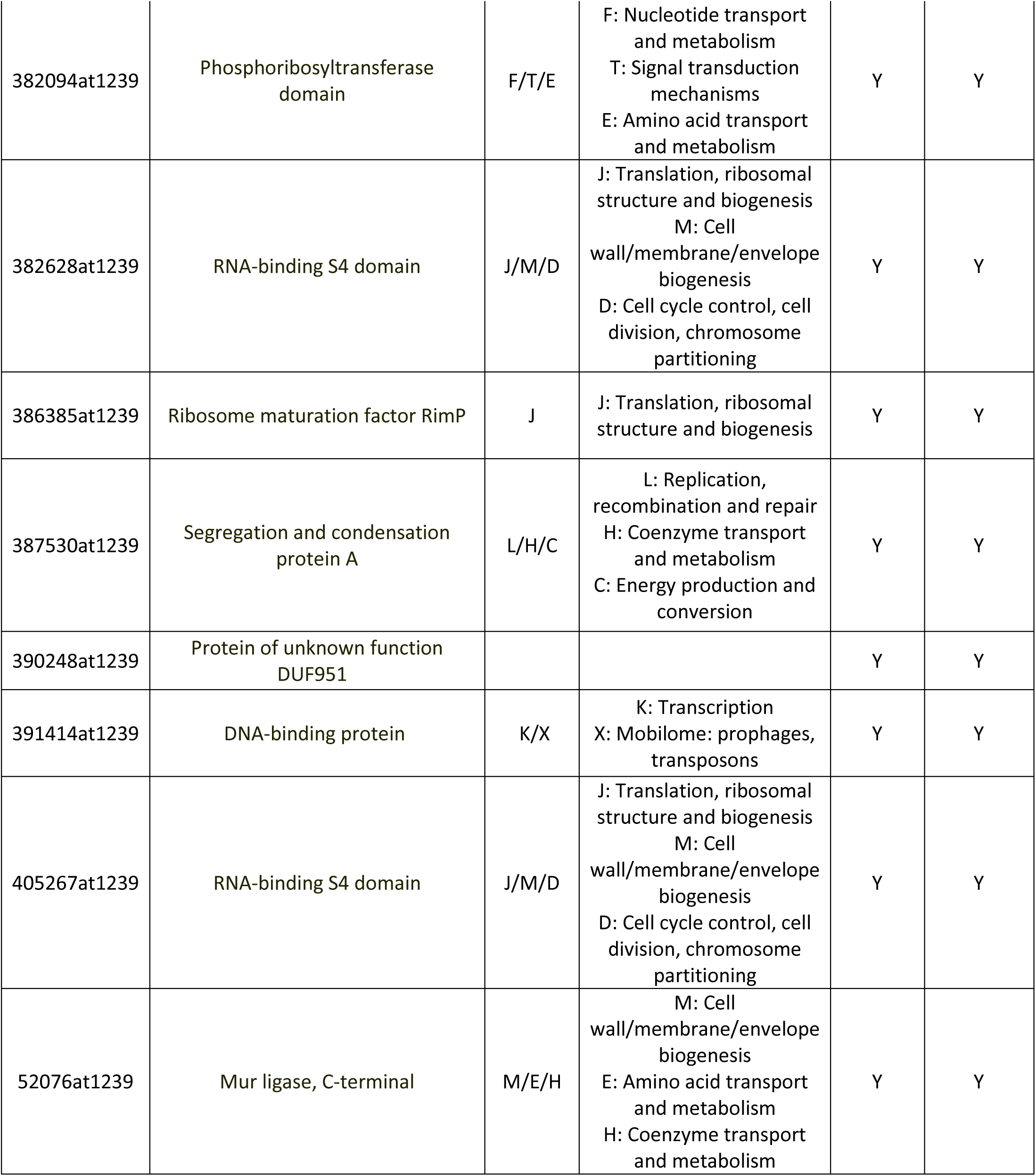
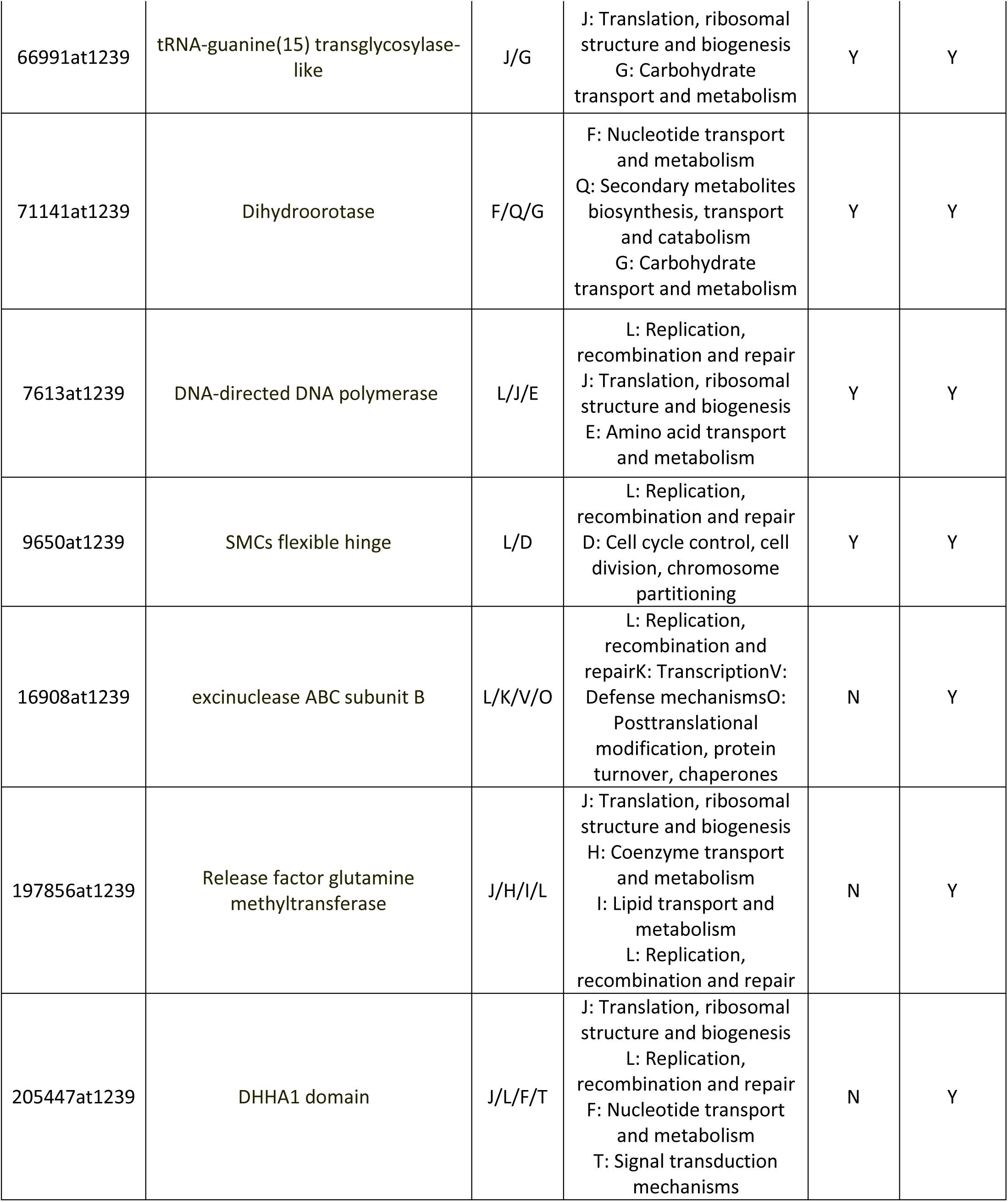
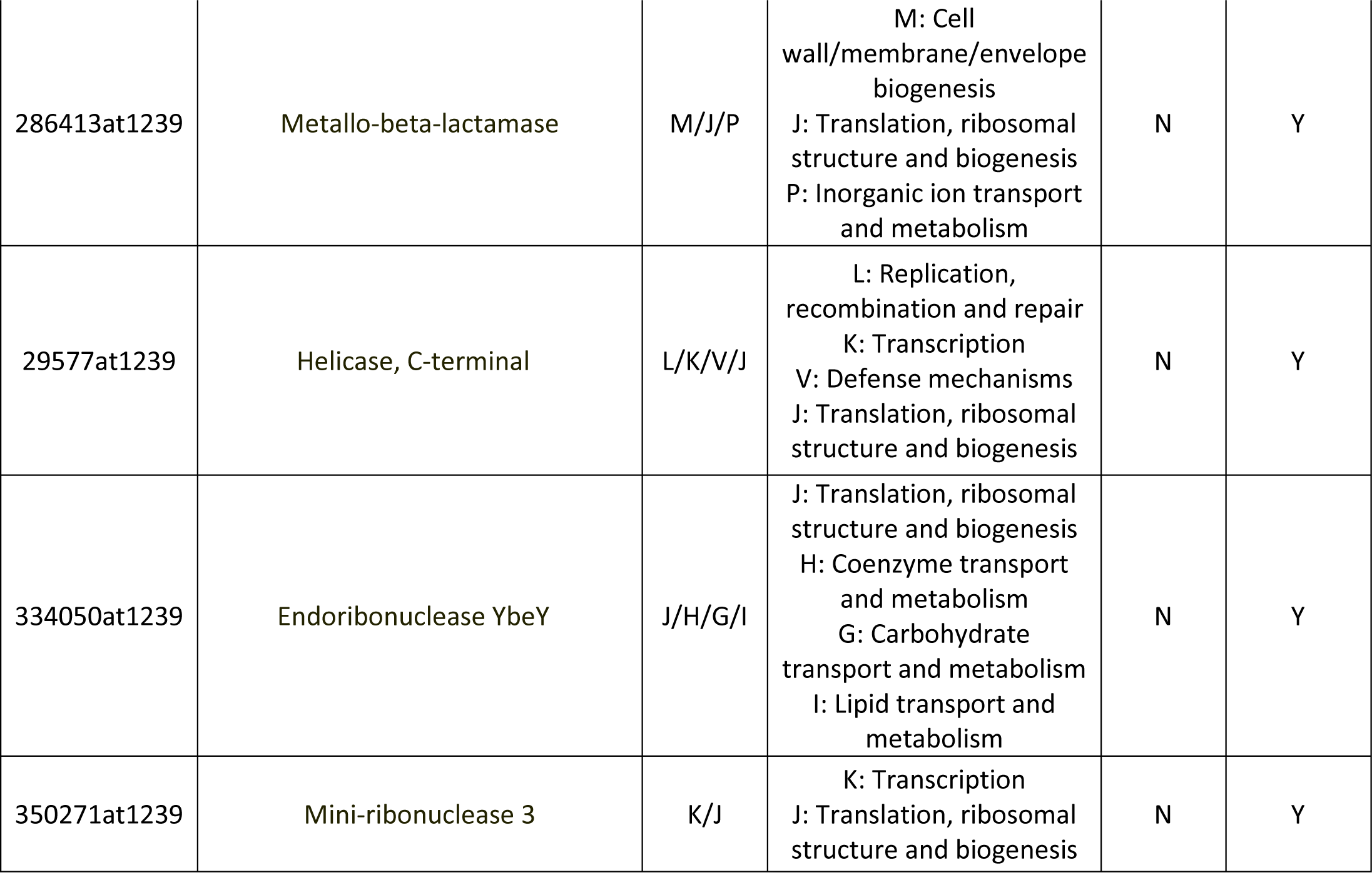
BUSCO genes missing in *P. ramosa* and functions.

**Table S3:**
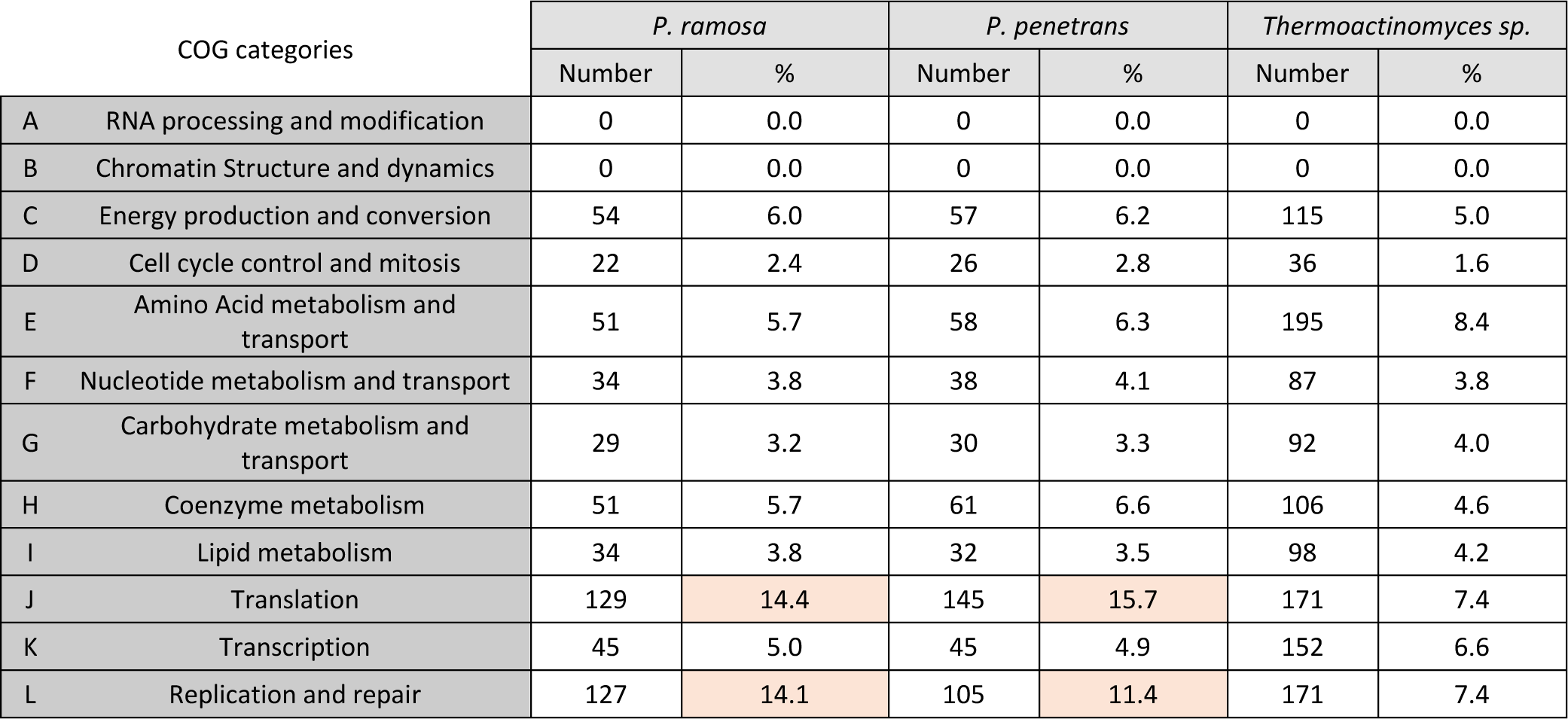
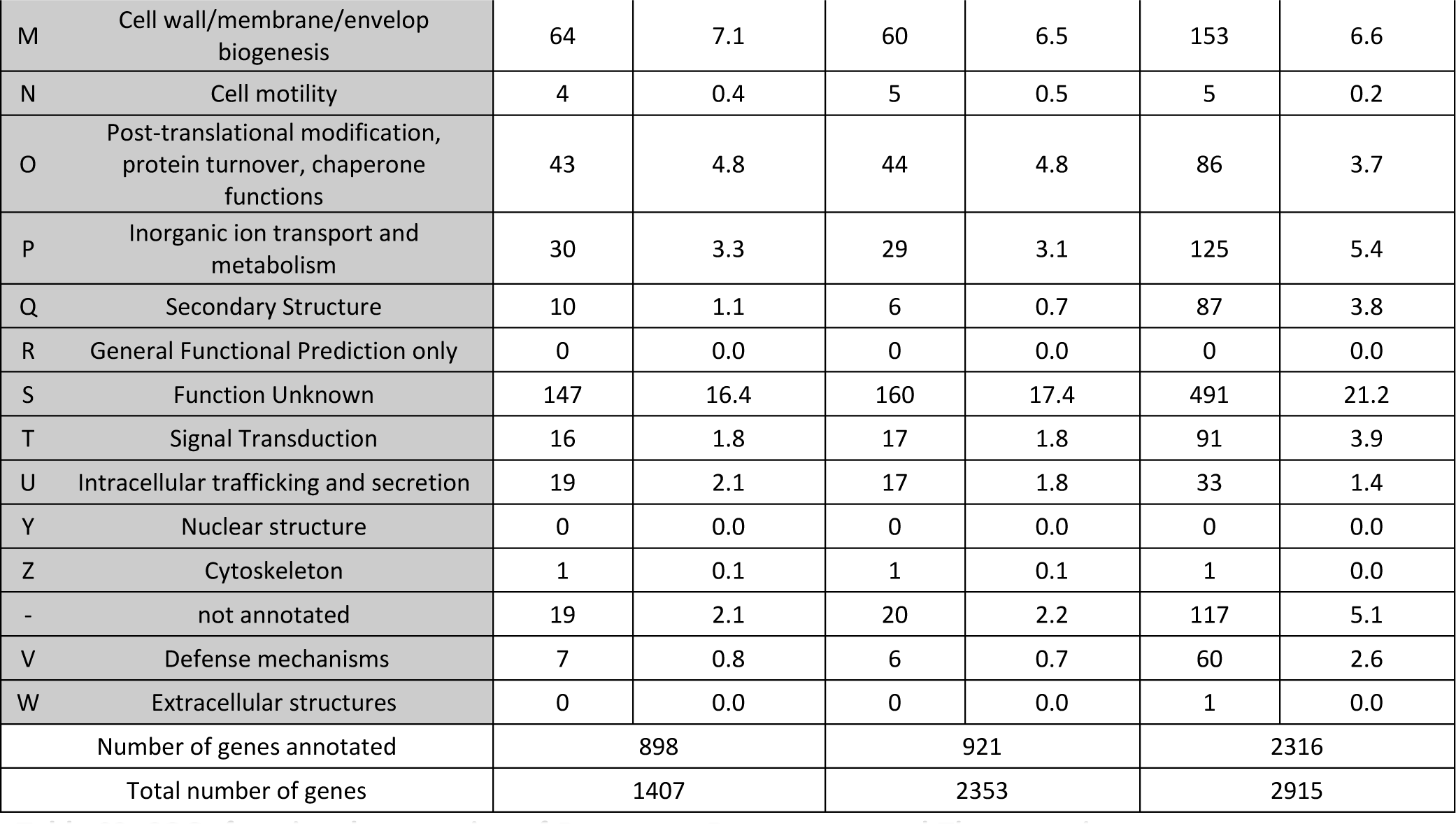
COGs functional annotation of *P. ramosa*, *P. penetrans* and *Thermoactinomyces sp.*. Percentage is the ratio between the annotation per categorie by the total number of proteins annotated.

**Table S4:** Genes in orthogroups specific to *P. ramosa*.

| Orthogroup | P. ramosa gene names |
| --- | --- |
| OG0000114 | NEKOFPGG_00066, NEKOFPGG_00068, NEKOFPGG_00069, NEKOFPGG_00072, NEKOFPGG_00073, NEKOFPGG_00187, NEKOFPGG_00195, NEKOFPGG_00230, NEKOFPGG_00231, NEKOFPGG_00232, NEKOFPGG_00233, NEKOFPGG_00235, NEKOFPGG_00240, NEKOFPGG_00241, NEKOFPGG_00250, NEKOFPGG_00251, NEKOFPGG_00606, NEKOFPGG_00607, NEKOFPGG_00624, NEKOFPGG_00625, NEKOFPGG_00642, NEKOFPGG_00682, NEKOFPGG_00683, NEKOFPGG_00784, NEKOFPGG_00823, NEKOFPGG_00851, NEKOFPGG_00852, NEKOFPGG_00867, NEKOFPGG_00976, NEKOFPGG_00993, NEKOFPGG_01050, NEKOFPGG_01081, NEKOFPGG_01082, NEKOFPGG_01083, NEKOFPGG_01084, NEKOFPGG_01144, NEKOFPGG_01145, NEKOFPGG_01191, NEKOFPGG_01216, NEKOFPGG_01228, NEKOFPGG_01229, NEKOFPGG_01230, NEKOFPGG_01249, NEKOFPGG_01259, NEKOFPGG_01313, NEKOFPGG_01314, NEKOFPGG_01316, NEKOFPGG_01318, NEKOFPGG_01368 |
| OG0001461 | NEKOFPGG_00258, NEKOFPGG_00260, NEKOFPGG_00366, NEKOFPGG_00645, NEKOFPGG_00680, NEKOFPGG_00791, NEKOFPGG_00822, NEKOFPGG_00893, NEKOFPGG_00895, NEKOFPGG_00907, NEKOFPGG_00995, NEKOFPGG_01181, NEKOFPGG_01182, NEKOFPGG_01384 |
| OG0001462 | NEKOFPGG_00259, NEKOFPGG_00261, NEKOFPGG_00364, NEKOFPGG_00365, NEKOFPGG_00576, NEKOFPGG_00577, NEKOFPGG_00790, NEKOFPGG_00821, NEKOFPGG_00892, NEKOFPGG_00894, NEKOFPGG_00906, NEKOFPGG_00994, NEKOFPGG_01183, NEKOFPGG_01383 |
| OG0002216 | NEKOFPGG_00227, NEKOFPGG_00900, NEKOFPGG_00968, NEKOFPGG_01124, NEKOFPGG_01171, NEKOFPGG_01192, NEKOFPGG_01245, NEKOFPGG_01260, NEKOFPGG_01307 |
| OG0002217 | NEKOFPGG_00753, NEKOFPGG_01107, NEKOFPGG_01110, NEKOFPGG_01111, NEKOFPGG_01112, NEKOFPGG_01114, NEKOFPGG_01115, NEKOFPGG_01116, NEKOFPGG_01119 |
| OG0002437 | NEKOFPGG_00228, NEKOFPGG_00899, NEKOFPGG_00967, NEKOFPGG_01125, NEKOFPGG_01189, NEKOFPGG_01246, NEKOFPGG_01261, NEKOFPGG_01306 |
| OG0003072 | NEKOFPGG_00542, NEKOFPGG_00543, NEKOFPGG_00544, NEKOFPGG_00545, NEKOFPGG_00546, NEKOFPGG_00547 |
| OG0003073 | NEKOFPGG_00910, NEKOFPGG_01298, NEKOFPGG_01299, NEKOFPGG_01300, NEKOFPGG_01301, NEKOFPGG_01302 |
| OG0003532 | NEKOFPGG_00052, NEKOFPGG_00975, NEKOFPGG_01016, NEKOFPGG_01080, NEKOFPGG_01175 |
| OG0004169 | NEKOFPGG_00054, NEKOFPGG_00055, NEKOFPGG_00138, NEKOFPGG_00785 |
| OG0004170 | NEKOFPGG_01176, NEKOFPGG_01286, NEKOFPGG_01370, NEKOFPGG_01371 |
| OG0005409 | NEKOFPGG_00578, NEKOFPGG_01242, NEKOFPGG_01382 |
| OG0005410 | NEKOFPGG_00746, NEKOFPGG_00747, NEKOFPGG_00748 |
| OG0005411 | NEKOFPGG_00811, NEKOFPGG_00812, NEKOFPGG_01162 |
| OG0005412 | NEKOFPGG_01027, NEKOFPGG_01028, NEKOFPGG_01029 |
| OG0005413 | NEKOFPGG_01070, NEKOFPGG_01075, NEKOFPGG_01204 |
| OG0007580 | NEKOFPGG_00159, NEKOFPGG_00160 |
| OG0007581 | NEKOFPGG_00242, NEKOFPGG_00589 |
| OG0007582 | NEKOFPGG_00412, NEKOFPGG_00414 |
| OG0007583 | NEKOFPGG_00456, NEKOFPGG_00457 |
| OG0007584 | NEKOFPGG_00590, NEKOFPGG_00591 |
| OG0007585 | NEKOFPGG_01126, NEKOFPGG_01247 |
| OG0007586 | NEKOFPGG_01177, NEKOFPGG_01178 |
| OG0007587 | NEKOFPGG_01386, NEKOFPGG_01389 |

### Files

File S1: Sequence of PCL 6-7-8 array 1 and 2

File S2: Methods Confirmation of the triplet PCL 6-7-8 duplication

